# Mechanical properties of DNA double-crossover motifs

**DOI:** 10.64898/2026.04.10.717625

**Authors:** Eva Matoušková, Marek Cuker, Filip Lankaš

## Abstract

DNA double-crossover (DX) molecules, comprising two Holliday junctions connected by two duplex arms, are fundamental building blocks of DNA nanostructures, but their mechanical properties remain poorly understood. Here we investigate the elasticity of isolated antiparallel DX motifs with 18 to 22 base pairs between the crossovers. Using mechanical models parameterized by extensive all-atom molecular dynamics simulations, we demonstrate that the bending rigidity of the duplexes within a DX motif is highly anisotropic, and that this anisotropy results from long-range elastic couplings involving all the duplex base pairs between the crossovers. The duplex stretch modulus decreases due to localized defects, while the twist stiffness is close to that of an isolated duplex. The DX core as a whole follows an analytical beam theory in bending but not in torsion. Our results extend beyond local elastic models of DNA nanostructures and pave the way for probing peculiar mechanical properties of other key motifs for DNA and RNA nanotechnology.

## Main text

DNA double-crossover (DX) molecules consist of two Holliday junctions connected by two double-helical domains (1). DX molecules can be parallel or antiparallel, depending on the relative orientation of the duplexes. The parallel molecules tend to dissociate or form multimers (1), therefore the main focus has been on the antiparallel ones. Among these, the antiparallel DX motif with even number of half-turns between the crossovers (DAE) comprises five strands, the central one being circular (1) (Figure 1). Early cyclization experiments (2) indicated that a DAE motif cyclizes no more readily than the linear molecule containing the same sequence, making it the long-sought unit to create periodic structures (3). The first higher-order DNA lattice structure, a two-dimensional crystal made of DX molecules, soon followed (4). Since then, DX motifs have been extensively employed to form densely packed DNA origami (5,6), edges of wireframe structures (7,8), or even photonic wires (9). Thus, DX motifs have become fundamental building blocks in the field of DNA nanotechnology.

**Figure 1.**
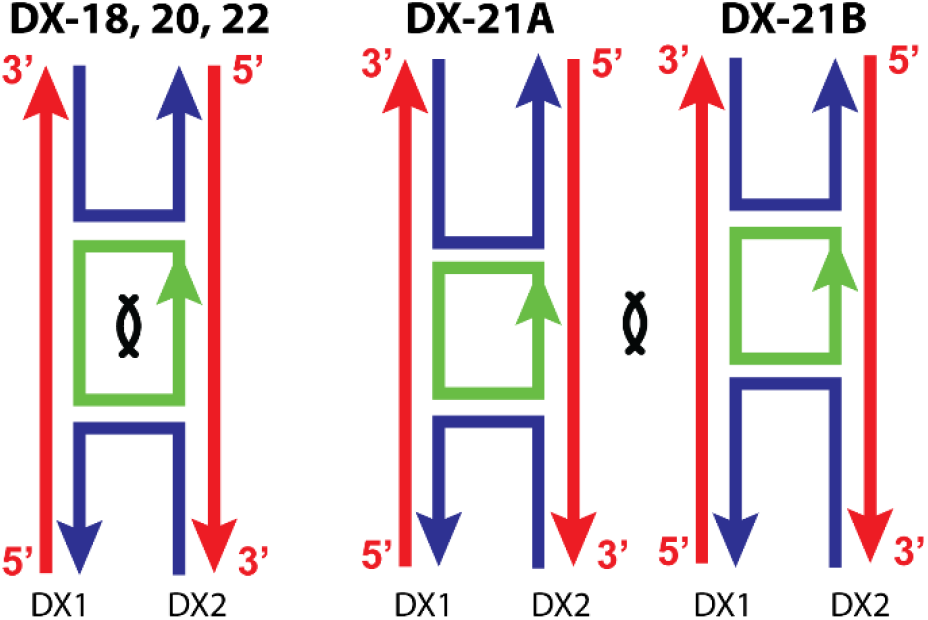
Double-crossover molecules DX-*N* investigated in this work, where *N* denotes the number of base pairs between the crossovers (within the circular strand). The two duplexes in each motif are denoted by DX1 and DX2. The two-fold symmetry, marked by the lens-like symbol, relates DX1 to DX2 in the same motif for even *N*, and DX1 in one to DX2 in the other molecule for *N* = 21.

DNA nanostructures are often subjected to mechanical stress, internal through scaffolding interactions inside the structure (10,11), which may also be introduced intentionally to create curved and twisted shapes (12,13), or external to harness the nanostructure compliance for functional purposes (14,15). The stress distribution within the structure, its conformational change, and stability all depend on the mechanical properties of the building blocks. For instance, the relative compliance of chained DX motifs (two-helix bundles, 2HB) compared to multihelix bundles was a key factor to realize mechanical frustration using DNA origami (16). Understanding the shape and stiffness of individual DX molecules is thus a prerequisite for assessing the structural response of these assemblies.

A major research effort has been devoted to the study of individual Holliday junctions (17-19). However, the conformation of a Holliday junction (HJ) in the DX molecule is very different from that of an isolated HJ (11,20). Moreover, the close proximity of the two junctions, or crossovers, may give rise to new mechanical phenomena. This implies the need to examine the DX motif, rather than an individual HJ, as a basic mechanical unit.

Experimental information about the structure and elasticity of DX molecules is surprisingly limited. The classical cyclization study (21), performed on constructs involving 3-8 chained DAE monomers, reported effective persistence length to be approximately twice as high as that of a linear duplex DNA. The stiffening was similar for the nicked and sealed circular strand. Only one duplex of the motif (the reporter domain) was cyclized in these experiments, while the other duplex was capped with a short loop at its ends. Thus, the data reflect the cyclizability of one duplex within the DX motif, rather than that of the motif as a whole. The axial stiffness of two-helix nanobeams made of DX motifs, with various densities of nicks and HJs, was examined by stretching in fluid flow (22). At the forces applied (*F* between 15 and 50 pN), the chains were surprisingly flexible with respect to stretching, a result consistent with the assumption of a local loss of stiffness at nicks and the crossover points. Flexible crossovers were also found in an origami rectangle folded into a cylindrical tube (23). These results indicate that parts of the DX motif may yield under force. Whether structural defects exist at zero force as well, and how they could possibly look like, is not known. Furthermore, stiffness defects outside the crossovers and nicks may exist as well, especially in the central part between the crossovers where the helices are severely bent (5,11). To our knowledge, the torsional rigidity of DX motifs has not been examined.

Computer simulations of DNA nanostructures have emerged as an indispensable complement to experimental investigations, providing insight into the structural dynamics in unprecedented detail, as well as design verification prior to time-consuming, costly experimental validation. The methods used to probe the mechanical properties of DNA assemblies range from all-atom molecular dynamics (MD) simulations (24,25), single-nucleotide models such as oxDNA (20,26), multiscale approaches (27), up to analytical (14,23,28) and finite-element models of continuum elasticity, of which CanDo (29) and SNUPI (30) are perhaps the most popular. None of these computational approaches has been applied to probe individual DX motifs. Recently, short (100 ns) all-atom MD simulations of isolated DX molecules have been reported, but the focus was not on their mechanical properties (31). Thus, little computational insight into the structure and elasticity of the DX molecules is available.

The continuum mechanics models (14,23,28-30) treat DNA nanostructures as assemblies of independent building blocks, be it a base-pair step within a duplex, or a crossover connecting two duplexes. Thus, the blocks are elastically uncoupled in these models and the total elastic energy is a sum of contributions from the individual blocks. However, it has long been understood that bases and base pairs in DNA duplexes are elastically coupled, resulting in length-dependent elasticity (32-36). A recent extensive study from our lab (36) demonstrated that elastic couplings span at least 5 base pairs (bp) along the DNA duplex and 4 bp along the RNA duplex. Whether significant elastic couplings exist in a DX molecule, and how they influence its mechanical properties, remains unknown.

In this study we present a mechanical characterization of isolated DX motifs in their relaxed state, i.e. at zero external force and torque. We focus on the most common antiparallel version with (roughly) even number of half-turns between the crossovers (DAE). At a coarser scale, we model the individual duplexes, and the whole domain between the crossovers (the DX core), as stretchable, twistable, anisotropically bendable elastic rods. We also explore a finer-grained description at the level of individual base pairs (bp), each treated as a rigid body. The models were parameterized from extensive atomic-resolution, explicit-solvent MD simulations. The duplex arms are 46 bp long. Each system comprises ca. 330,000 atoms and was simulated for 5 µs, providing both locally and globally converged model parameters. An isolated DNA duplex was simulated as a control.

A series of molecules with *N* inter-crossover pairs (*N* = 18, 20, 21, 22, two versions for *N* = 21), denoted DX-18, DX-20, DX-21A, DX-21B and DX-22, were investigated. The sequences of the continuous strands in the two duplex domains are the same (Figure S1) and the circular strand is sealed, so that the molecules with even *N* enjoy an exact two-fold symmetry, while the two *N* = 21 structures were designed so that they are connected by a two-fold symmetry to each other (Figure 1). The structures were built manually using PyMol, and the Amber suite of programs was used for the simulations, employing the OL15 DNA force field (37), 150 mM KCl added salt (38), the SPC/E water model (39), and a standard simulation protocol (36). The first 200 ns were excluded from the analysis due to initial large structural rearrangements. From that point on, the molecules were stable, with only short, reversible breaks of non-terminal pairs.

To characterize a duplex fragment as a flexible rod, we define four internal, global (G) coordinates. The global twist (G-twist) is computed as a sum of helical twists of the individual base-pair steps, the fragment length is a sum of helical rises. Helical twists and rises are calculated as in the 3DNA conformational analysis program (40). This description, already proved useful to investigate duplex mechanics (36,41,42), is extended here by introducing the global roll and global tilt angles (G-roll, G-tilt) to capture the anisotropic bending. To do so, we define one coordinate frame at each end of the fragment, for which we take the 3DNA base-pair frame at that location. We further choose a frame in the center of the fragment, namely the 3DNA bp step frame of the duplex central step (Figure S2). The mean normal of the two end frames defines a middle plane, onto which the x-axis of the central frame is projected, defining the x-direction of a middle frame (Figure S3). The rest of the definition is exactly the same as for the local roll and local tilt in 3DNA (40). Thus, G-roll quantifies bending out of the plane of the DX core, whereas G-tilt captures the in-plane bending. The global coordinates were measured on each snapshot of the analyzed MD trajectories. The stretch modulus *Y* was obtained from the variance of the duplex length, the twist stiffness *C* from the variance of the global twist, and the twist-stretch coupling *k* by inverting the 2-by-2 covariance matrix of the two, as in (36,41). Similarly, inverting the 2-by-2 covariance matrix of G-roll and G-tilt gives the out-of-plane bending stiffness *A*_*ro*_, the in-plane bending stiffness *A*_*ti*_, and the coupling term *A*_*roti*_ (Table S1).

Figures 2 and S4 show the elastic constants for duplex fragments starting at the central bp step and elongated symmetrically by one step at each side. While the out-of-plane bending stiffness *A*_*ro*_ is only slightly higher than that of an isolated duplex, the in-plane bending stiffness *A*_*ti*_ is about three times larger at the crossovers, and is still nearly twice as high 7 bp away from the crossovers, compared to the free duplex.

**Figure 2.**
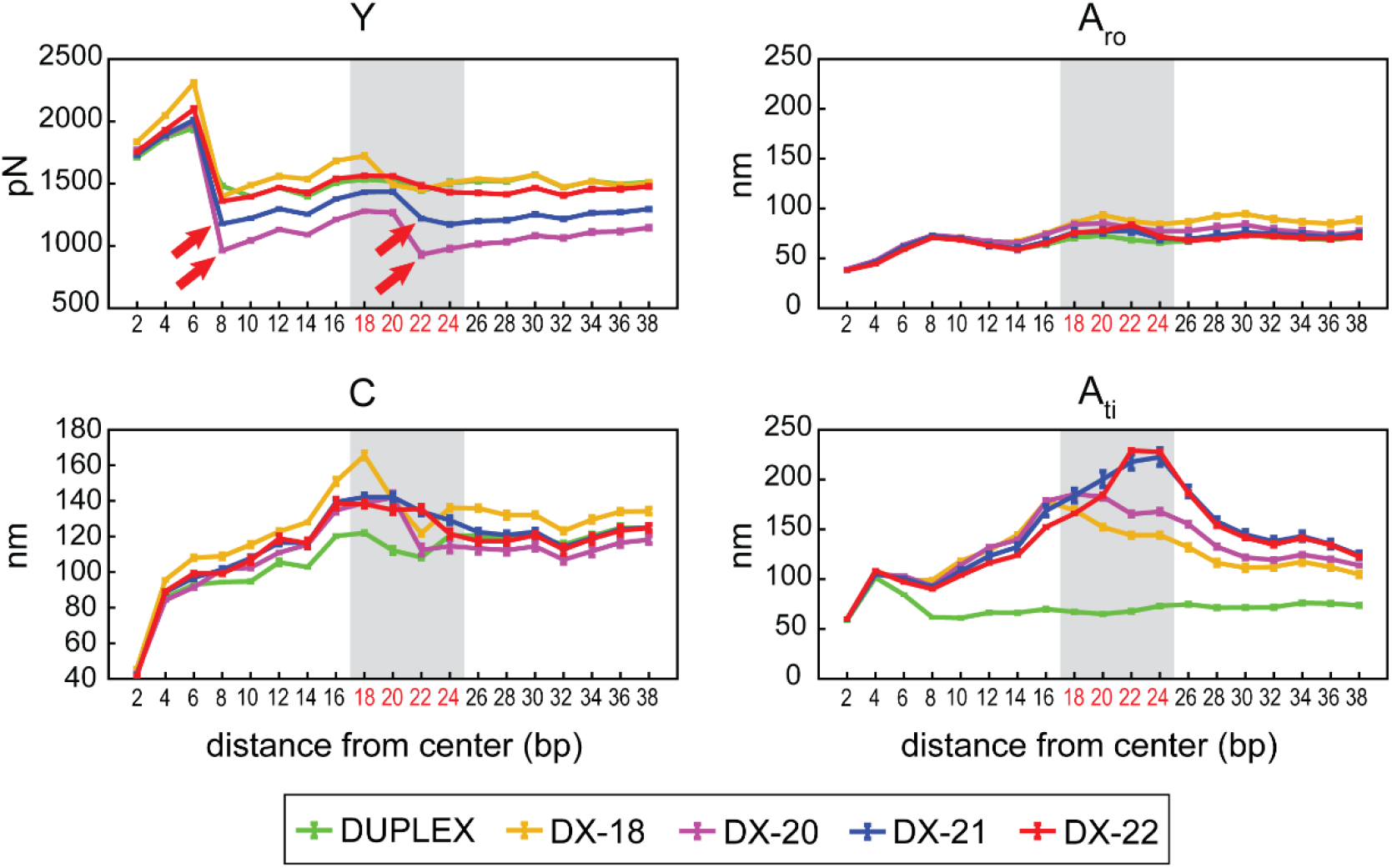
Elastic constants for duplexes in DX motifs and for the isolated duplex inferred from MD simulation data. The fragments start at the duplex center and are elongated symmetrically on both sides towards the ends. The domain where the crossovers are located is marked in grey. The locations of structural defects are highlighted by red arrows. The values for the two symmetry-related duplexes (Figure 1) are very close to each other and are averaged in the plots. Average errors for the two duplexes, each computed as the absolute difference between values for the whole trajectory and for its halves, are also shown.

Thus, mounting a DNA duplex into a DX motif profoundly changes the duplex bending elasticity well beyond the crossover points.

These results may have important ramifications for the mechanics of DNA nanostructures. In particular, they provide a mechanistic understanding for the intriguing cyclization data on chained DX motifs (21), which indicate a two-fold increase in bending rigidity of DX-mounted duplexes compared to isolated ones and which, > 20 years after publication, remain unexplained. The cyclization method cannot resolve the bending anisotropy and only reports an effective isotropic value of the duplex bending stiffness. One may take the bending stiffness *A*_*iso*_, the harmonic mean of the two eigenvalues of the 2-by-2 bending stiffness matrix (43), as an approximation of the measured bending stiffness. We see that *A*_*iso*_ of a duplex in the DX motif is already 40-70 % higher than that of an isolated duplex at the crossovers, and still visibly higher 7 bp away (Figure S4). However, how exactly the cyclization proceeds is not entirely clear, as the other, capped duplexes may present a significant steric hindrance. We hypothesize that the in-plane bending may be somewhat preferred for steric reasons, since then the capped duplexes would be at the outer side of the circle. Our data, suggesting that the in-plane bending is stiff, would then imply an even larger effective bending stiffness than *A*_*iso*_, in line with the experiment.

To get insight into the underlying mechanism of the bending anisotropy, we described each duplex at a finer-grained level, namely that of rigid base pairs. The configuration of the duplex of *m* base pairs is given by 6(*m* − 1) base-pair step coordinates (6 for each step: shift, slide, rise, tilt, roll and twist), for which we again used the 3DNA definitions. In addition to the raw MD data, we generated an ensemble of configurations based on the MD-derived coordinate means and coordinate covariance matrix ∑, assuming a normal distribution (a Monte Carlo method), and analyzed the structures as if they were snapshots from an MD trajectory. If the full covariance matrix ∑ was used, the resulting anisotropic bending stiffness was nearly indistinguishable from the original MD data (Figure 3A, B). If, by contrast, just a block diagonal version of ∑ was employed, neglecting couplings between the 6-by-6 blocks representing individual steps, the anisotropy was completely lost and, moreover, the duplex was markedly more flexible (Figure 3B). This demonstrates that the bending anisotropy of DNA duplexes within a DX motif is due to elastic couplings between individual base-pair steps in the duplex.

**Figure 3.**
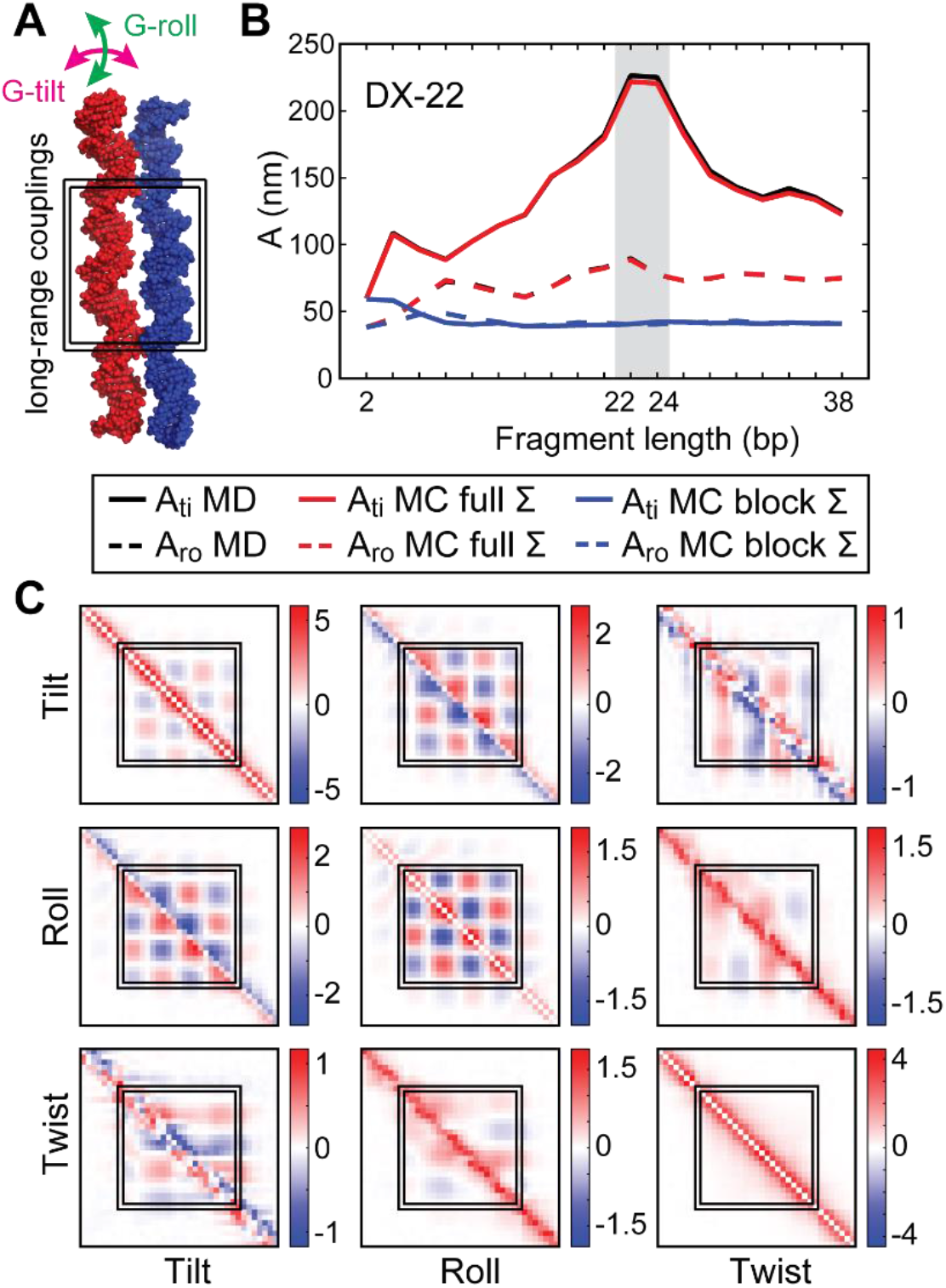
The nature of long-range elastic couplings in the DX motif. Data for DX-22 are shown as an example. (A) A representative structure and a schematic definition of the two bending angles. (B) The bending stiffness constants from raw MD data, from an MD-informed normal distribution with the full base-pair step covariance matrix ∑, and from a normal distribution with the block-diagonal ∑ where elastic couplings between different steps are set to zero. (C) Entries of the full stiffness matrix corresponding to rotational coordinates. See Figures S5-S9 for the other motifs.

To further investigate the nature of these couplings, we computed the full stiffness matrix *K* = *k*_*B*_ *T* ∑^−1^, normalized it using a length scale of 1 Å and angle scale of 11° (36), and re-arranged its entries so that the coordinates of the same type (twist, roll and so on) are grouped together. While the translational (shift-shift, shift-slide etc.), or the translational-rotational mixed entries (shift-tilt etc.) showed only a limited coupling range reminiscent of an isolated duplex, the rotational coordinates were coupled all over the section of the duplex between the crossovers, but not any further (Figures 3C and S5). Thus, the decisive couplings to induce the bending anisotropy involve local rotational coordinates (tilt, roll, twist) between the crossovers.

A recent study from our lab (36) indicated that the elastic couplings between bases in a DNA or RNA duplex are predominantly not of electrostatic origin, since they remain nearly unchanged if the duplex is artificially neutralized in the simulations. Instead, the mechanism may be analogous to the “railway-track model” (44) whereby two worm-like chains connected by harmonic springs exhibit a higher bending stiffness. Similar ideas were used to explain length-dependent stiffness of a DNA duplex (35,45). Given that tightly packed DNA origami exhibit intricately woven DNA backbones, the non-local coupling is likely to be widespread in these structures. In particular, our results suggest that duplexes in single layered DNA origami (5,6) are likely extremely rigid for in-plane bending. Long-range elastic couplings may also affect wireframe structures where the edges are formed by duplexes rather than DX tiles (46). The nature and effective range of the coupling may depend on the boundary conditions, i.e. the way the duplexes are anchored.

Turning now to the duplex stretching stiffness, we observe values similar to those for the isolated duplex, but also significant dips at local structural defects (Figure 2). These appear in DX-20 and DX-21 molecules where the helices are close together, whereas the more open DX-18 and DX-22 are defect-free. Apart from the crossovers, there are defects also in the central part, presumably due to the sharp bending of the duplexes between the crossovers. The atomistic simulations enable us to characterize these structural anomalies in detail. They consist of local buckling of the double helix, exhibiting small helical rise, high inclination of the base pairs with respect to the helical axis, anomalous values of backbone dihedral angles, and high variance of the helical rise (Figures 4 and S10), which results in a local decrease of the stretch modulus. By contrast, the twist stiffness, apart from a local rigidification at the crossovers, remains close to that of an isolated duplex (Figure 2). Finally, the twist-stretch coupling is somewhat reduced (Figure S4). We verified that the stiffness constants for the symmetry-related duplexes are very close to each other, indicating excellent global convergence of our simulated systems (Figures S6-S9).

**Figure 4.**
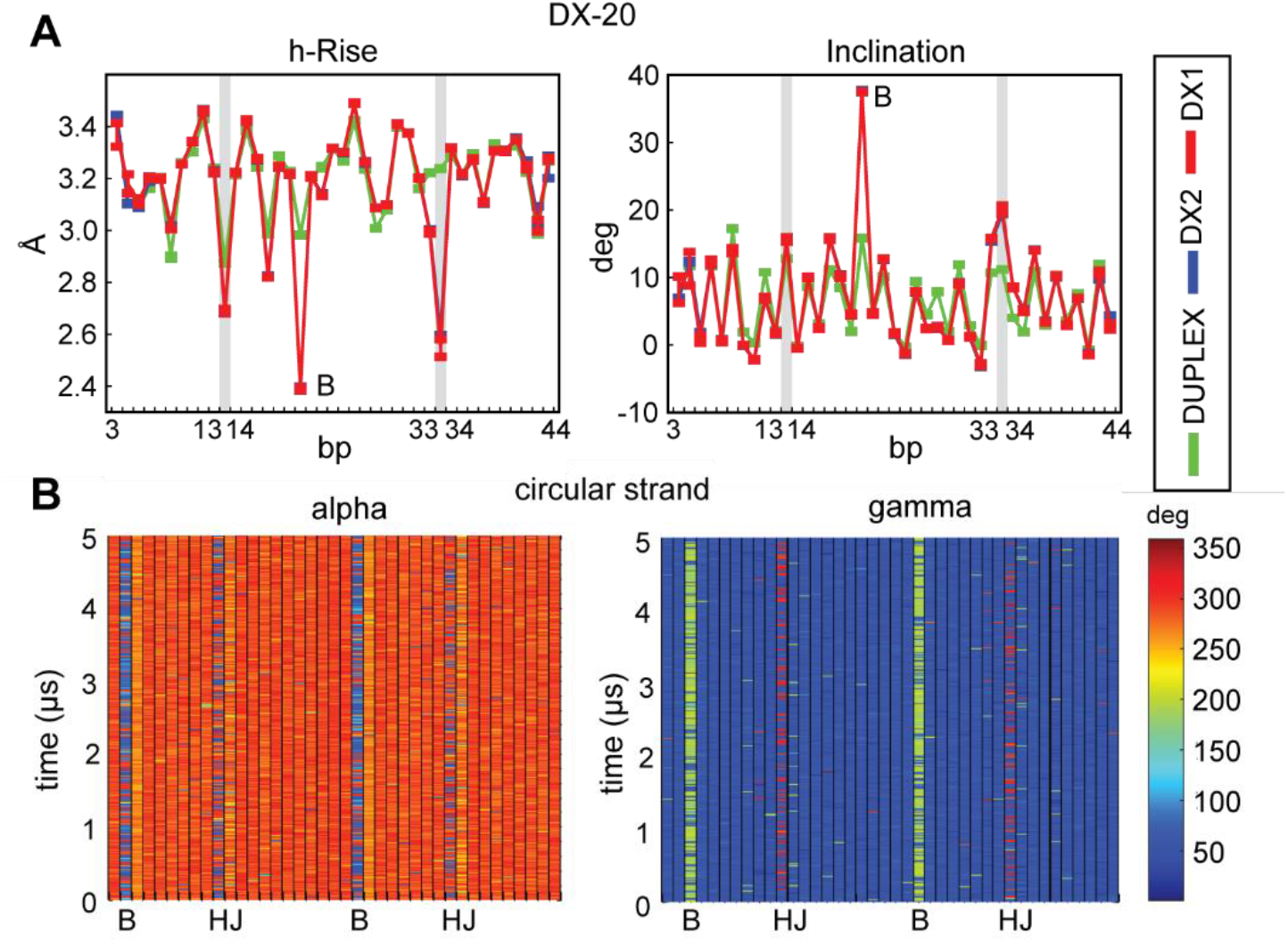
Structural defects in DX-20. (A) Helical rise and inclination for the two duplexes in the motif (red and blue). Data for the isolated duplex (green) are shown for comparison. Defects are found within the Holliday junctions (“HJ”, grey stripes), and between the junctions (“B”) where the helix is locally buckled. (B) Time series of the backbone torsion angles α and γ.

Just as the individual duplexes within the DX motif, we also model the whole DX core as a flexible rod. Here the base-pair step frames in the four crossover steps each represent a virtual base, the adjacent crossover steps in the two duplexes a virtual pair, and the core a virtual, giant base-pair step (Figure S11). We define the core bending angles in exact analogy to the 3DNA roll and tilt angles, while we take the core length as the average length, and the core twist as the average G-twist (i.e. sum of helical twists) of the two duplexes.

If the duplexes were straight, homogeneous, isotropic rods with circular cross-section, then according to the Euler-Bernoulli (EB) theory of elastic beams, *A*_*ro*_ of the core should be twice the value of an isolated duplex and *A*_*ti*_ 10 times that value, so that the ratio of *A*_*ti*_ to *A*_*ro*_ should be equal to 5 (SI Methods). The inferred bending stiffnesses of the core indeed follow these rules rather well, with the exception of the open DX-18 motif (Table S2). Thus, while the individual duplexes within the DX motif substantially differ from an isolated duplex in that they are highly anisotropic, the core as a whole is well approximated in bending as a pair of isotropic DNA duplexes. This is likely due to the geometric effect of the shifted neutral axis which dominates the in-plane bending stiffness of the core (SI Methods), and which would also explain the success of the EB theory to model bending of other tightly packed, homogeneous helical bundles (14,28,47). By contrast, we expect the long-range elastic couplings to play a major role in assemblies where the stiffness of individual, interconnected duplexes dominates.

The stretching stiffness of the core inferred here is, as expected, roughly twice the value of a single duplex (Tables S1 and S2). As for the torsional stiffness, we are unaware of any analytical beam theory for the double-circular cross-section representing the DX core. We therefore approximated the cross-section with a rectangle into which the two circles are inscribed (Figure S12). Then a standard theory of torsion for rectangular beams predicts that the torsional stiffness of the beam should be 4.67 that of the individual duplex (SI Methods). However, our DX core is just about twice as stiff in torsion as an isolated duplex (Tables S1 and S2). This is also in line with magnetic tweezer experiments on four-helix and six-helix bundles, where the twist stiffness of the bundle was found roughly proportional to the number of duplexes in the bundle, while finite-element-model (FEM) simulations of interconnected duplexes predicted a significantly higher torsional stiffness (47).

Taken together, in this work we proposed structural models at various levels of detail and parameterized them from extensive all-atom MD simulations to deduce the mechanical properties of isolated double-crossover motifs. Our results indicate that all the base pairs of a duplex within the DX core are elastically coupled, challenging standard local models of DNA nanostructure elasticity. The coupling results in highly anisotropic bending stiffness of the duplexes in the motif. This suggests a mechanistic basis for experimental cyclization data and exposes the potential role of long-range couplings in other structures built from interconnected duplexes. The duplex stretching stiffness may decrease due to local defects in and outside the crossovers, here characterized in atomistic detail, while the twist stiffness remains similar to that of an isolated duplex. We also establish the elastic properties of the DX core as a whole and discuss them in light of theoretical predictions for elastic beams. Our work demonstrates the utility of a multiscale approach to uncover intricate mechanical features of a key DNA nanostructure motif, and may be readily extended to model mechanical properties of other building blocks for DNA and RNA nanotechnology.

## Supporting information

SI methods (theory of elastic beams), SI figures showing coordinate definitions and additional results, SI tables with numerical values of the elastic constants (pdf).

## Acknowledgments

This work was supported by Specific University Research at the University of Chemistry and Technology Prague [A1_FCHT_2024_001 to E.M.].

## Table of contents graphics

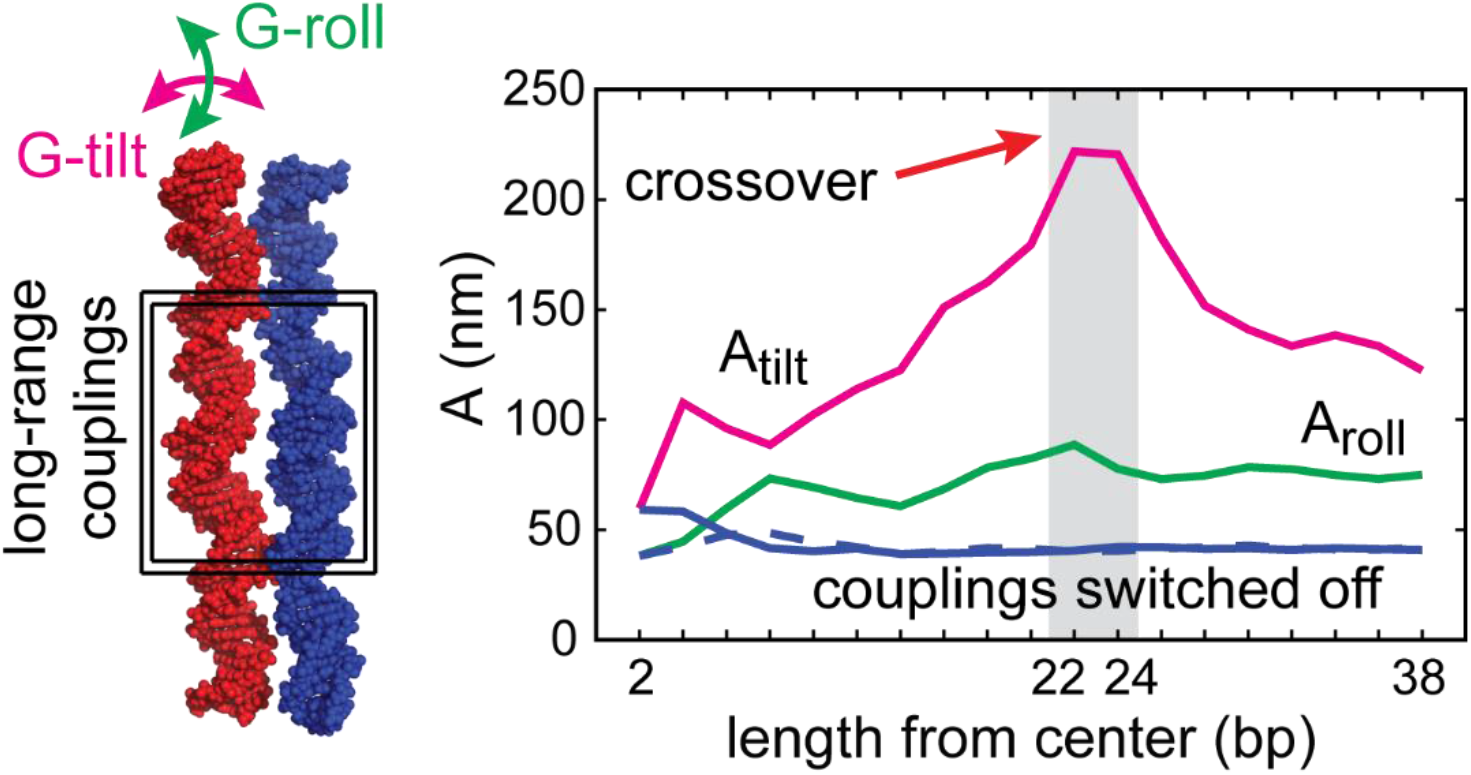

## References

1. Fu, T.-J. and Seeman, N.C. (1993) DNA double-crossover molecules. Biochemistry, 32, 3211–3220.

2. Li, X., Yang, X., Qi, J. and Seeman, N.C. (1996) Antiparallel DNA double crossover molecules as components for nanoconstruction. J. Am. Chem. Soc., 118, 6131–6140.

3. Seeman, N.C. (1998) DNA nanotechnology: novel DNA constructions. Annu. Rev. Biophys. Biomol. Struct., 27, 225–248.

4. Winfree, E., Liu, F., Wenzler, L.A. and Seeman, N.C. (1998) Design and self-assembly of two-dimensional DNA crystals. Nature, 394, 539–544.

5. Rothemund, P.W.K. (2006) Folding DNA to create nanoscale shapes and patterns. Nature, 440, 297–302.

6. Hong, F., Zhang, F., Liu, Y. and Yan, H. (2017) DNA origami: Scaffolds for creating higher order structures. Chem. Rev., 117, 12584–12640.

7. Zhang, F., Jiang, S., Wu, S., Li, Y., Mao, C., Liu, Y. and Yan, H. (2015) Complex wireframe DNA origami nanostructures with multi-arm junction vertices. Nat. Nanotechnol., 10, 779–784.

8. Veneziano, R., Ratanalert, S., Zhang, K., Zhang, F., Yan, H., Chiu, W. and Bathe, M. (2016) Designer nanoscale DNA assemblies programmed from the top down. Science, 352, aaf4388.

9. Scott, M.N., Banal, J.L., Chen, W.J., Brooks, C., Wang, X., Hart, S.M., Dodin, A., Bathe, M., Willard, A.P. and Schlau-Cohen, G.S. (2025) Transport of delocalized excitons through DNA-based molecular photonic wires. ACS Nano, 19, 38509–38520.

10. Ke, Y., Douglas, S.M., Liu, M., Sharma, J., Cheng, A., Leung, A., Liu, Y., Shih, W.M. and Yan, H. (2009) Multilayer DNA origami packed on a square lattice. J. Am. Chem. Soc., 131, 15903–15908.

11. Bai, X.-C., Martin, T.G., Scheres, S.H.W. and Dietz, H. (2012) Cryo-EM structure of a 3D DNA-origami object. Proc. Natl. Acad. Sci. USA, 109, 20012–20017.

12. Dietz, H., Douglas, S.M. and Shih, W.M. (2009) Folding DNA into twisted and curved nanoscale shapes. Science, 325, 725–730.

13. Han, D., Pal, S., Nangreave, J., Deng, Z., Liu, Y. and Yan, H. (2011) DNA origami with complex curvatures in three-dimensional space. Science, 332, 342–346.

14. Zhou, L., Marras, A.E., Su, H.-J. and Castro, C.E. (2014) DNA origami compliant nanostructures with tunable mechanical properties. ACS Nano, 8, 27–34.

15. Li, R., Madhvacharyula, A.S., Du, Y., Adepu, H.K. and Choi, J.H. (2023) Mechanics of dynamic and deformable DNA nanostructures. Chem. Sci., 14, 8018–8046.

16. Madhvacharyula, A.S., Li, R., Swett, A.A., Du, Y., Seo, S., Simmel, F.C. and Choi, J.H. (2025) Realizing mechanical frustration at the nanoscale using DNA origami. Nature Commun., 16, 5164.

17. Lilley, D.M.J. (2000) Structures of helical junctions in nucleic acids. Q. Rev. Biophys., 33, 109–159.

18. Addendorf, M.R., Tang, G.Q., Millar, D.P., Bathe, M. and Bricker, W.P. (2021) Computational investigation of the impact of core sequence on immobile DNA four-way junction structure and dynamics. Nucleic Acids Res., 50, 717–730.

19. Simmons, C.R., MacCulloch, T., Krepl, M., Matthies, M., Buchberger, A., Crawford, I., Sponer, J., Sulc, P., Stephanopoulos, N. and Yan, H. (2022) The influence of Holliday junction sequence and dynamics on DNA crystal self-assembly. Nature Commun., 13, 3112.

20. Snodin, B.E., Schreck, J.S., Romano, F., Louis, A.A. and Doye, J.P.K. (2019) Coarse-grained modelling of the structural properties of DNA origami. Nucleic Acids Res., 47, 1585–1597.

21. Sa-Ardyen, P., Vologodskii, A. and Seeman, N.C. (2003) The flexibility of DNA double crossover molecules. Biophys. J., 84, 3829–3837.

22. Jung, W.-H., Chen, E., Veneziano, R., Gaitanaros, S. and Chen, Y. (2020) Stretching DNA origami: effect of nicks and Hollliday junctions on the axial stiffness. Nucleic Acids Res., 48, 12407–12414.

23. Chen, H., Weng, T.-W., Riccitelli, M.M., Cui, Y., Irudayaraj, J. and Choi, J.H. (2014) Understanding the mechanical properties of DNA origami tiles and controlling the kinetics of their folding and unfolding reconfiguration. J. Am. Chem. Soc., 136, 6995–7005.

24. Yoo, J. and Aksimentiev, A. (2013) In situ structure and dynamics of DNA origami determined through molecular dynamics simulations. Proc. Natl. Acad. Sci. USA, 110, 20099–20104.

25. Pandey, L., Panigaj, M., Radwan, Y., Chhabra, H., Chen, Y., Aksimentiev, A., Afonin, K.A. and Wanunu, M. (2025) Chemical composition and backbone modifications define deformability of nucleic acid nanoparticles. ACS Nano, 19, 24972–24984.

26. Chhabra, H., Mishra, G., Cao, Y., Presern, D., Skoruppa, E., Tortora, M.M.C. and Doye, J.P.K. (2020) Computing the elastic mechanical properties of rodlike DNA nanostructures. J. Chem. Theory Comput., 16, 7748–7763.

27. Maffeo, C. and Aksimentiev, A. (2020) MrDNA: a multi-resolution model for predicting the structure and dynamics of DNA systems. Nucleic Acids Res., 48, 5135–5146.

28. Schiffels, D., Liedl, T. and Fygenson, D.K. (2013) Nanoscale structure and microscale stiffness of DNA nanotubes. ACS Nano, 8, 6700–6710.

29. Pan, K., Bricker, W.P., Ratanalert, S. and Bathe, M. (2017) Structure and conformational dynamics of scaffolded DNA origami nanoparticles. Nucleic Acids Res., 45, 6284–6298.

30. Park, J.H., Kim, D.-N. and Lee, J.Y. (2025) SNUPI: a computational framework for rapid mechanical analysis of structured DNA assemblies. JACS Au, 5, 5813–5820.

31. Rodriguez, A., Madhanagopal, B.R., Sarkar, K., Nowzari, Z., Mathivanan, J., Talbot, H., Patel, A., Morya, V., Halvorsen, K., Vangaveti, S. et al. (2025) Counterions influence the isothermal self-assembly of DNA nanostructures. Sci. Adv., 11, eadu7366.

32. Lankas, F., Sponer, J., Hobza, P. and Langowski, J. (2000) Sequence-dependent elastic properties of DNA. J. Mol. Biol., 299, 695–709.

33. Noy, A. and Golestanian, R. (2012) Length scale dependence of DNA mechanical properties. Phys. Rev. Lett., 109, 228101.

34. Skoruppa, E., Voorspoels, A., Vreede, J. and Carlon, E. (2021) Length-scale-dependent elasticity in DNA from coarse-grained and all-atom models. Phys. Rev. E, 103, 042408.

35. Fosado, Y.A.G., Landuzzi, F. and Sakaue, T. (2023) Coarse graining DNA: Symmetry, nolocal elasticity, and persistence length. Phys. Rev. Lett., 130, 058402.

36. Slavnikova, P., Cuker, M., Matouskova, E., Cmelo, I., Zgarbova, M., Jurecka, P. and Lankas, F. (2025) Sequence-dependent shape and stiffness of DNA and RNA double helices: hexanucleotide scale and beyond. J. Chem. Inf. Model., 65, 9208–9229.

37. Zgarbova, M., Sponer, J., Otyepka, M., Cheatham III, T.E., Galindo-Murillo, R. and Jurecka, P. (2015) Refinement of the sugar-phosphate backbone torsion beta for Amber force fields improves the description of Z- and B-DNA. J. Chem. Theory Comput., 11, 5723–5736.

38. Dang, L.X. (1995) Mechanism and thermodynamics of ion selectivity in aqueous solutions of 18-crown-6 ether: a molecular dynamics study. J. Am. Chem. Soc., 117, 6954–6960.

39. Berendsen, H.J.C., Grigera, J.R. and Straatsma, T.P. (1987) The missing term in effective pair potentials. J. Phys. Chem., 91, 6269–6271.

40. Lu, X.-J. and Olson, W.K. (2003) 3DNA: a software package for the analysis, rebuilding and visualization of three-dimensional nucleic acid structures. Nucleic Acids Res., 31, 5108–5121.

41. Liebl, K., Drsata, T., Lankas, F., Lipfert, J. and Zacharias, M. (2015) Explaining the striking difference in twist-stretch coupling between DNA and RNA: A comparative molecular dynamics analysis. Nucleic Acids Res., 43, 10143–10156.

42. Marin-Gonzalez, A., Vihena, J.G., Perez, R. and Moreno-Herrero, F. (2017) Understanding the mechanical response of double-stranded DNA and RNA under constant stretching force using all-atom molecular dynamics. Proc. Natl. Acad. Sci. USA, 114, 7049–7054.

43. Landau, L.D. and Lifshitz, E.M. (1980) Statistical Physics, Part 1. Elsevier, Amsterdam.

44. Everaers, R., Bundschuh, R. and Kremer, K. (1995) Fluctuations and stiffness of double-stranded polymers: Railway-rack model. Europhys. Lett., 29, 263–268.

45. Segers, M., Voorspoels, A., Sakaue, T. and Carlon, E. (2022) Mechanical properties of nucleic acids and the non-local twistable wormlike chain model. J. Chem. Phys., 156, 234105.

46. Benson, E., Mohammed, A., Rayneau-Kirkhope, D., Gadin, A., Orponen, P. and Hogberg, B. (2018) Effects of design choices on the stiffness of wireframe DNA origami structures. ACS Nano, 12, 9291–9299.

47. Kauert, D.J., Kurth, T., Liedl, T. and Seidel, R. (2011) Direct mechanical measurements reveal the material properties of three-dimensional DNA origami. Nano Lett., 11, 5558–5563.

