## Supporting information for "Mechanical properties of DNA double-crossover motifs"

### Supporting methods

According to the Euler-Bernoulli theory of elastic beams (1,2), the bending stiffness $\tilde{A}$ of a straight, homogeneous, isotropic beam is given by $\tilde{A} = EI$, where $E$ is the Young's modulus (or modulus of elasticity) of the material and $I$ is the second moment of area of the beam's cross-section, calculated with respect to the axis perpendicular to the applied load (the neutral axis). For a beam of a circular cross-section of radius $r$, we have

$$I_c = \frac{1}{4}\pi r^4$$

for any bending direction. If the beam consists of two circular rods adjacent to each other (Figure S12), then the in-plane (tilt) and out-of-plane (roll) bending stiffnesses will be different. We obtain

$$I_{ro} = 2 \cdot \frac{1}{4}\pi r^4 = \frac{1}{2}\pi r^4$$

and, using the parallel axis theorem,

$$I_{ti} = 2 \cdot \left(\frac{1}{4}\pi r^4 + \pi r^2 \cdot r^2\right) = \frac{5}{2}\pi r^4$$

so that $\tilde{A}_{ti}/\tilde{A}_{ro} = I_{ti}/I_{ro} = 5$. Notice that the second term in the parentheses, appearing due to the shift of the neutral axis from the duplex center to the midpoint between the two duplexes, dominates the second moment of area and therefore also the in-plane bending stiffness. It is common to express the bending stiffness in units of length by introducing the quantity $A = \tilde{A}/k_B T$, where $k_B$ is the Boltzmann constant and $T$ the thermodynamic temperature.

The torsional stiffness $\tilde{C}$ of a straight, homogeneous, isotropic beam of a circular cross-section is given by $\tilde{C}_c = GJ_c$, where $G$ is the modulus of rigidity (or shear modulus of elasticity) of the material and $J_c$ the polar moment of inertia,

$$J_c = \frac{1}{2}\pi r^4$$

For beams of rectangular cross-section, a standard approximation (1) yields $\tilde{C}_r = GJ_r$, where for a rectangle of sides $a \geq b$,

$$J_r = c_2 a b^3$$

and $c_2$ is a coefficient depending on the ratio $a/b$ of the side lengths. In our case of two adjacent cylinders of radius $r$ approximated by a rectangle (Figure S12), we have $a = 4r$, $b = 2r$, $a/b = 2$ and $c_2 = 0.229$ (1). Thus, $\tilde{C}_r/\tilde{C}_c = J_r/J_c = 4.67$. It is common to express the torsional stiffness in units of length by introducing the quantity $C = \tilde{C}/k_B T$.

### Supporting figures

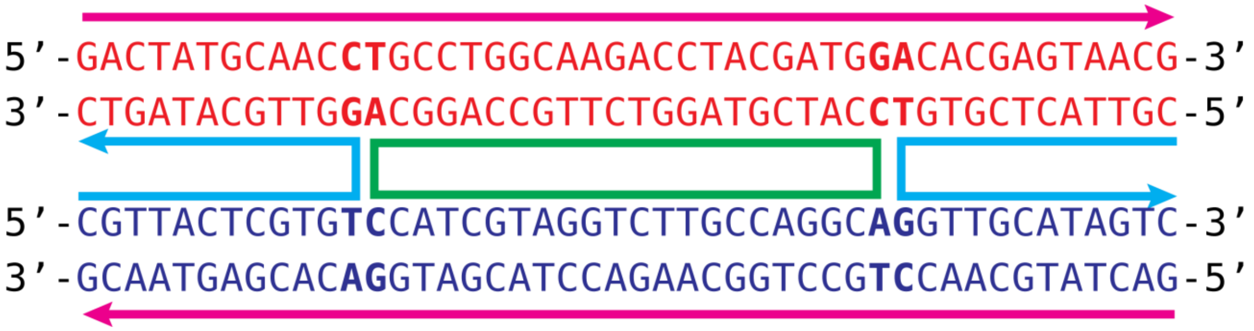


**Figure S1.** Sequence of the DX molecules investigated in this work. The duplexes have 46 bp each. The position of the crossovers shown corresponds to the DX-22 molecule. To construct the DX-20 and DX-18 motifs, the crossovers were shifted symmetrically on both sides towards the center. To obtain the DX-21A and DX-21B constructs, one crossover was shifted, while the other remained in place. An isolated duplex of the same sequence was also simulated as a control.

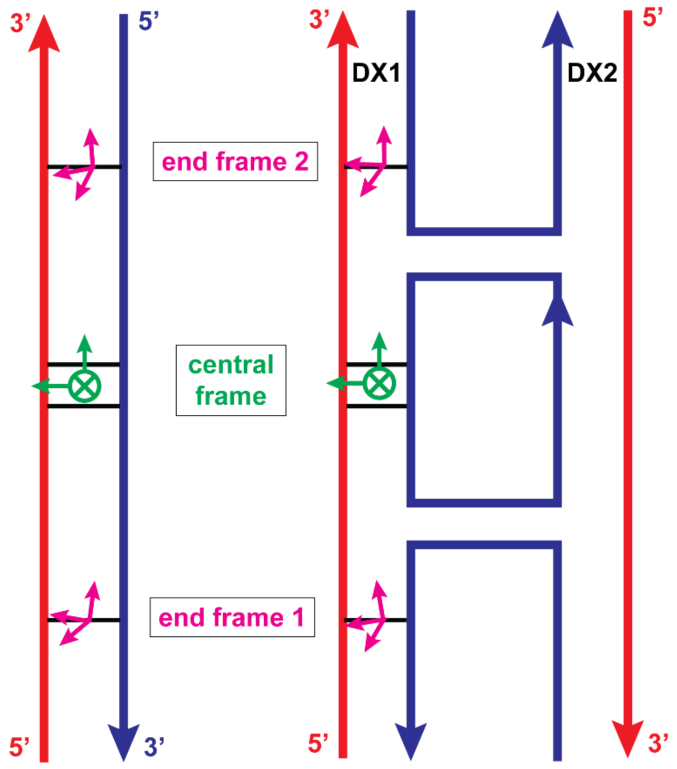


**Figure S2.** Frames used to define the global (G) coordinates for the duplex, either isolated (left) or within a DX molecule (right). The end frames (magenta) are base-pair frames of the pairs at the ends of the fragment, the central frame (green) is the bp step frame of the middle step of the fragment. The x-axis of the central frame, $\mathbf{x}_c$, points into the major groove (away from the viewer), the y-axis points to the left and the z-axis upwards in the figure. Thus, $\mathbf{x}_c$ in the DX molecules is perpendicular to the DX core in its center.

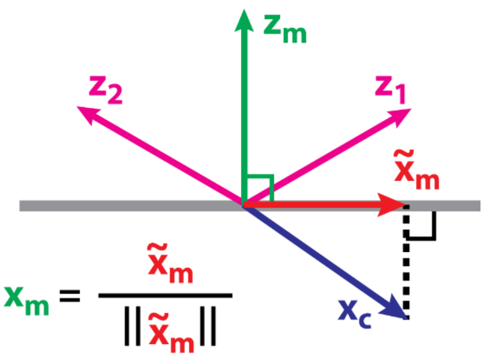


**Figure S3.** Definition of the middle frame for the duplex. The mean normal of the end frames defines the z-axis of the middle frame. The x-axis of the central frame (Fig. 2) is projected onto the plane perpendicular to the mean normal, defining the x-axis of the middle frame, while its y-axis complements the right-handed triad. Once the middle frame is defined, the coordinates are computed exactly as in 3DNA (3).

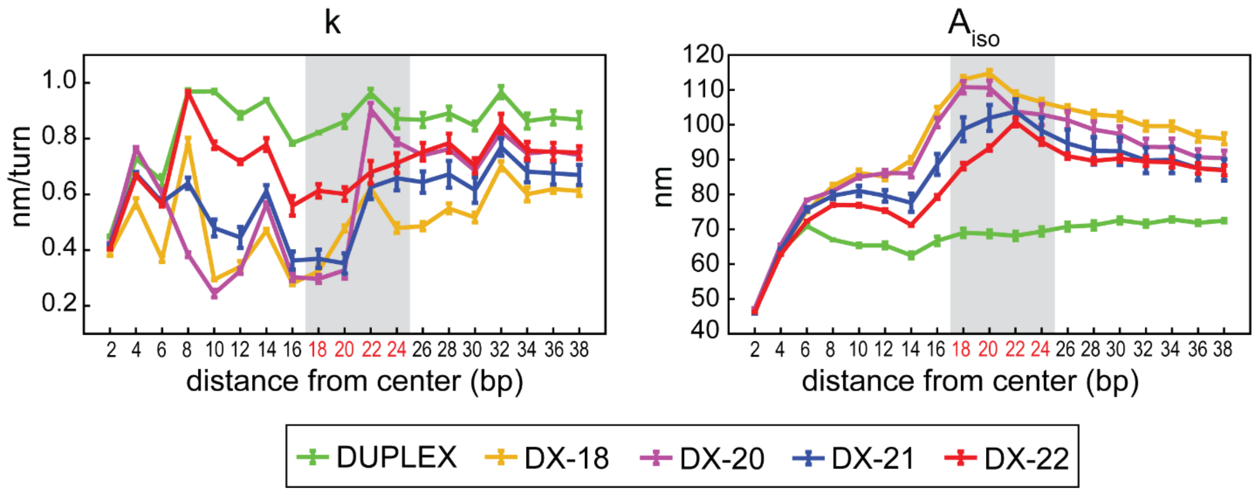


**Figure S4.** The twist-stretch coupling $k$ and the effective isotropic bending stiffness $A_{iso}$.

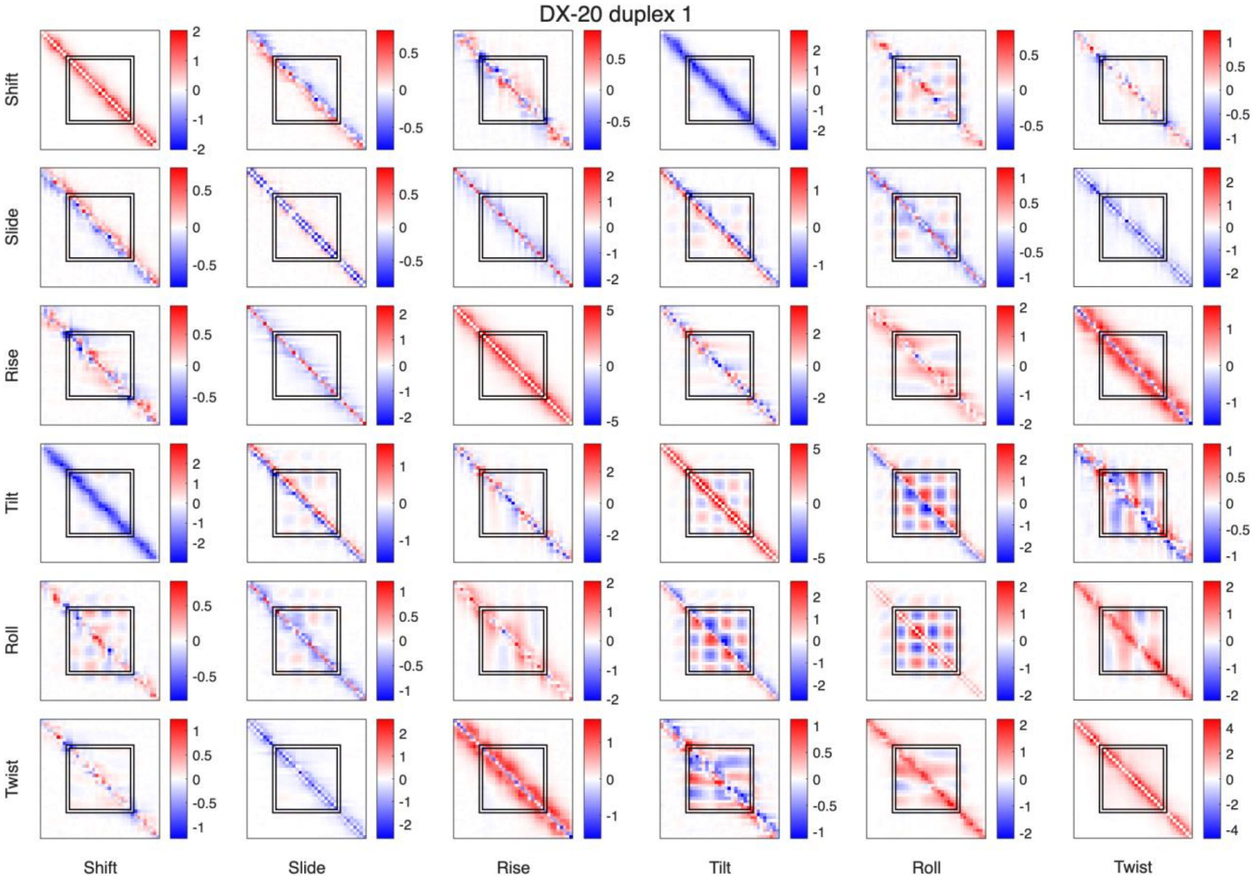


**Figure S5.** The stiffness matrix $K$ for one duplex in the DX-20 motif. Data for the other duplex in the same motif, and for duplexes in the other motifs, are entirely analogous.

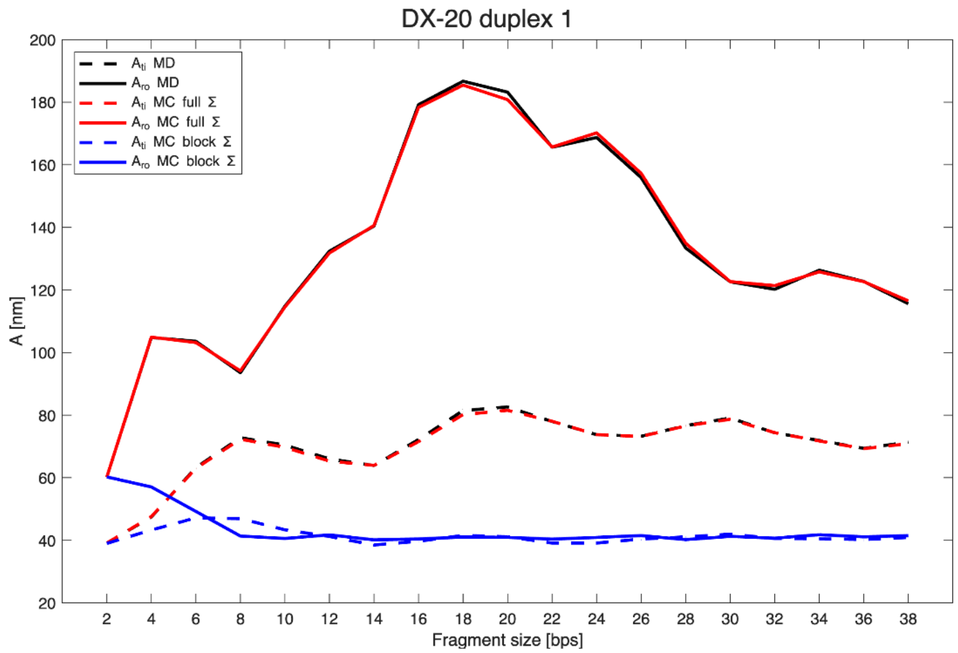


**Figure S6.** Raw MD and Monte Carlo (MC) generated bending stiffness data for one duplex of the DX-20 motif.

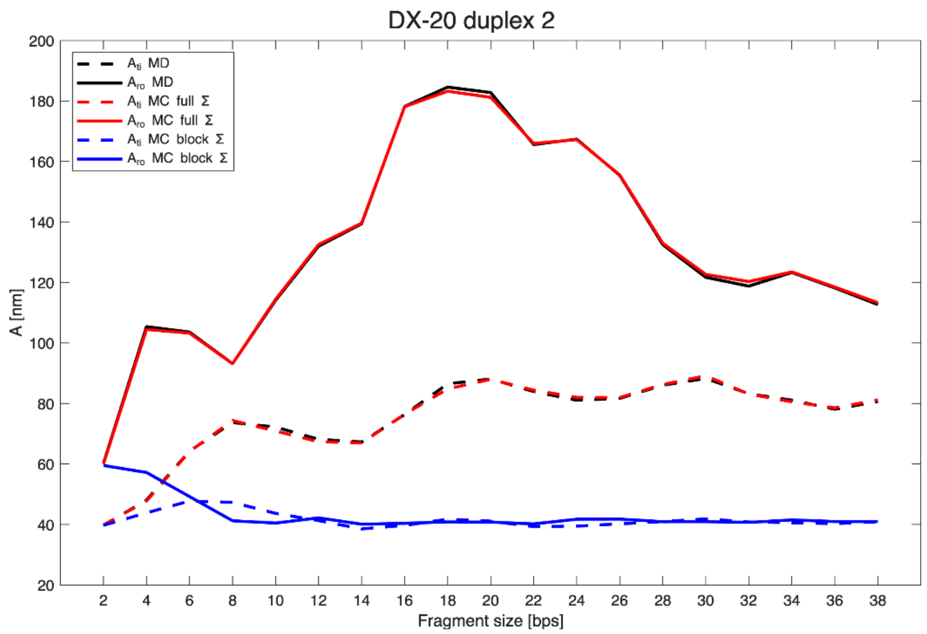


**Figure S7.** Raw MD and Monte Carlo (MC) generated bending stiffness data the other duplex of the DX-20 motif. The data are nearly identical with those in Fig. S6, indicating excellent global convergence.

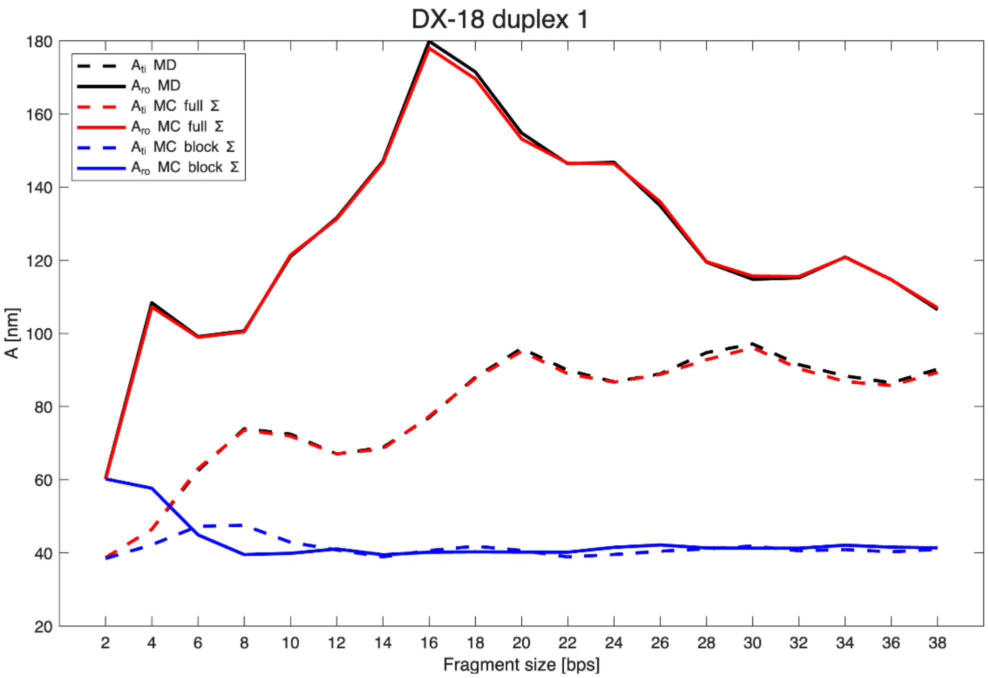


**Figure S8.** Raw MD and Monte Carlo (MC) generated bending stiffness data for one duplex of the DX-18 motif.

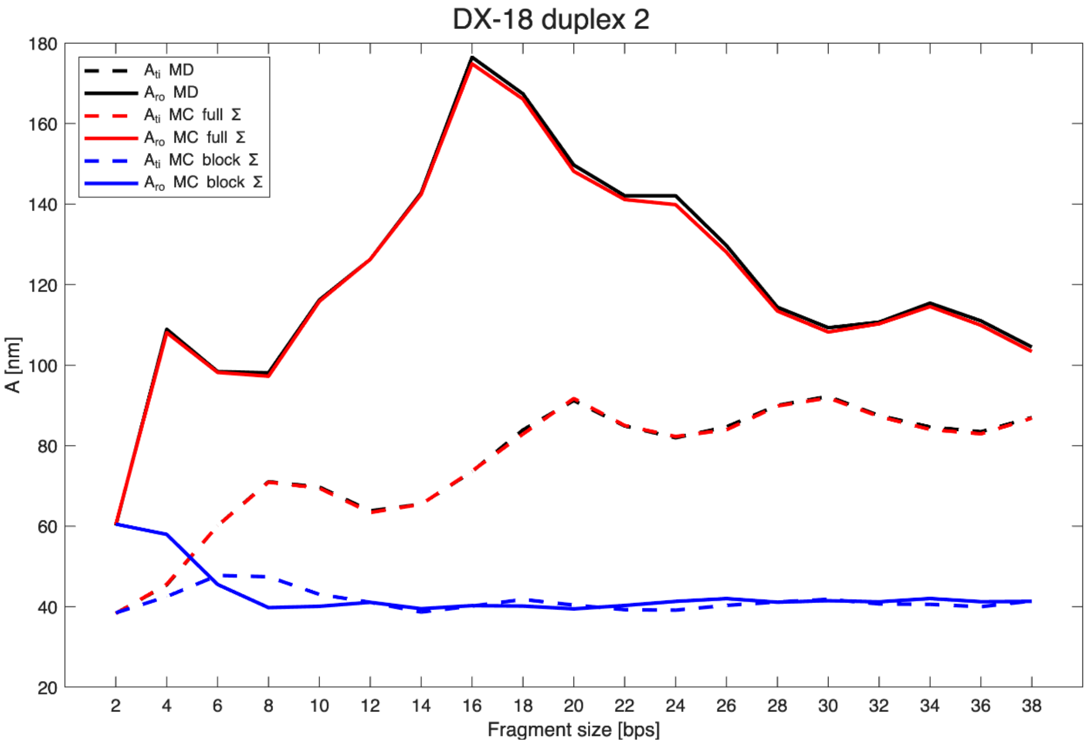


**Figure S9.** Raw MD and Monte Carlo (MC) generated bending stiffness data for the other duplex of the DX-18 motif.

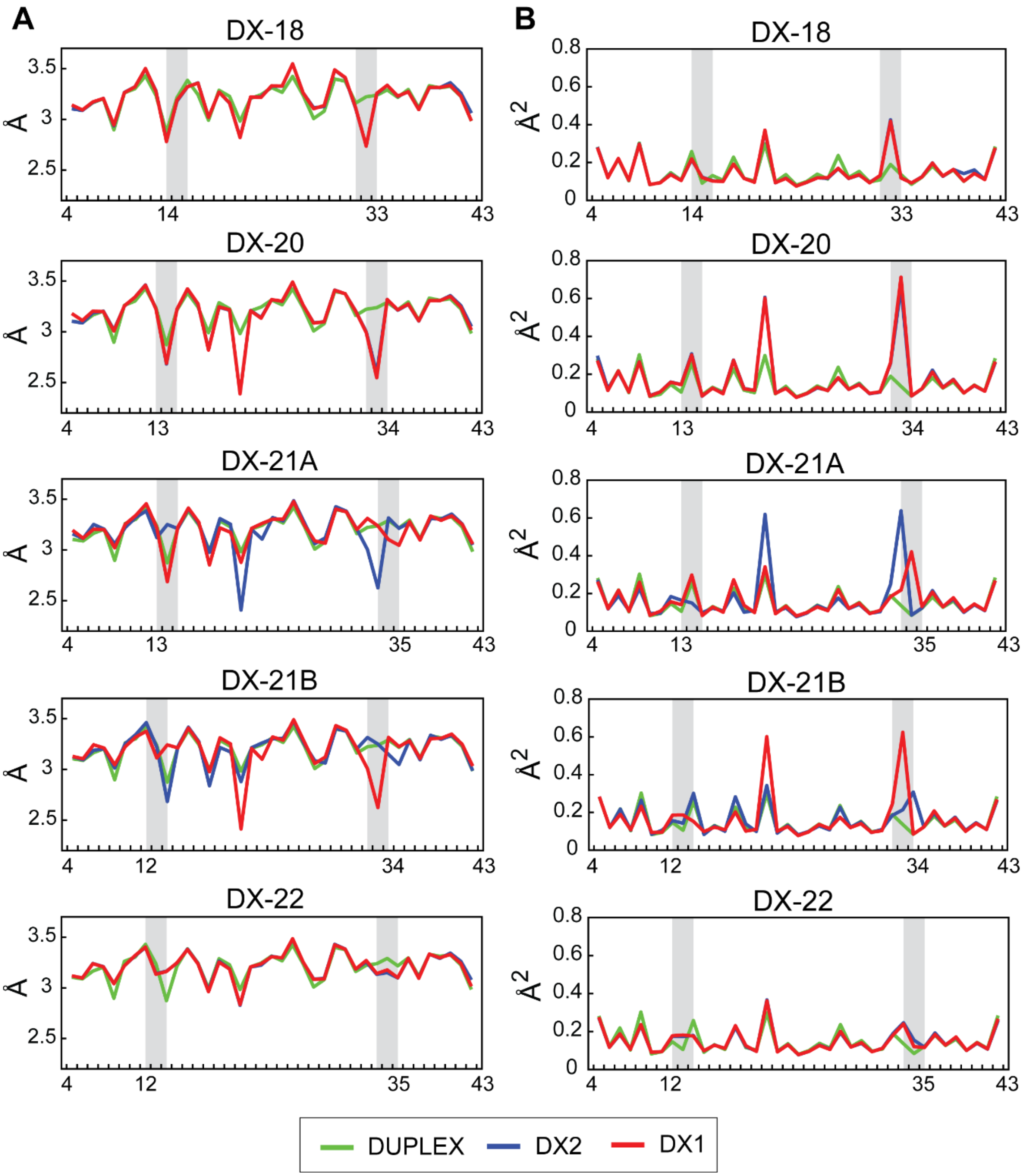


**Figure S10.** Mean and variance of the helical rise along the duplexes in DX motifs and the isolated duplex. Clear anomalies are seen in one duplex of DX-21A, B, and in both duplexes in DX-20, while DX-18 and DX-22 are defect-free. Notice the nearly identical values at the corresponding positions in symmetry-related duplexes. This indicates excellent convergence of the MD data.

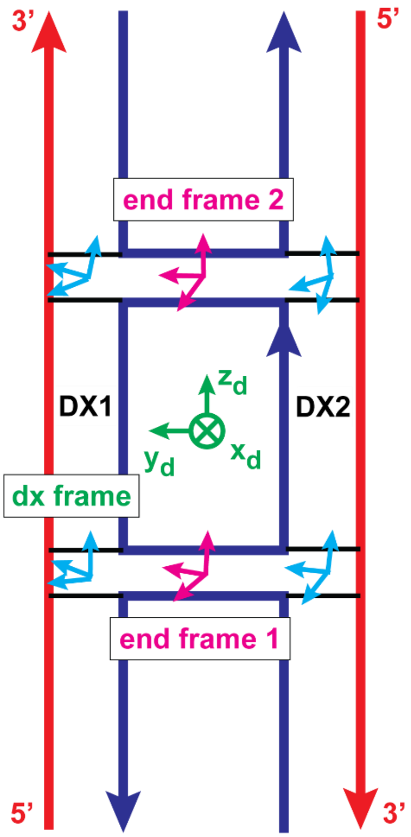


**Figure S11.** Definition of frames for the DX core. The middle frame (or dx frame) is computed from the end frames exactly as in the 3DNA algorithm.

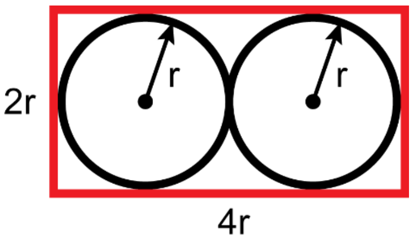


**Figure S12.** The DX core cross-section was approximated by a rectangle to estimate its twist stiffness.

### Supplementary tables

**Table S1.** Stiffness constants for duplexes in DX motifs (roman font) between the outer pairs of the crossovers, compared to the corresponding fragments of the isolated duplex (*italics*). Values for the individual duplexes are averaged. Mean errors are also shown.

| | **n = 18** | **n = 20** | **n = 21** | **n = 22** |
|---|---|---|---|---|
| **Y (pN)** | 1490.78 ± 7.14<br>*1520.41 ± 0.38* | 930.10 ± 16.11<br>*1441.25 ± 2.39* | 1172.60 ± 9.25<br>*1507.34 ± 18.04* | 1431.62 ± 8.66<br>*1507.34 ± 18.04* |
| **C (nm)** | 140.76 ± 1.04<br>*112.11 ± 1.73* | 112.38 ± 2.89<br>*108.28 ± 1.01* | 129.15 ± 2.10<br>*120.55 ± 2.14* | 121.47 ± 2.41<br>*120.55 ± 2.14* |
| **k (nm/turn)** | 0.48 ± 0.01<br>*0.86 ± 0.02* | 0.91 ± 0.02<br>*0.96 ± 0.01* | 0.66 ± 0.04<br>*0.87 ± 0.03* | 0.71 ± 0.04<br>*0.87 ± 0.03* |
| **$A_{ro}$ (nm)** | 93.55 ± 2.04<br>*72.84 ± 1.43* | 80.91 ± 3.59<br>*68.59 ± 1.21* | 70.14 ± 1.96<br>*65.98 ± 1.29* | 71.91 ± 1.09<br>*65.98 ± 1.29* |
| **$A_{ti}$ (nm)** | 152.20 ± 2.96<br>*65.13 ± 0.74* | 165.41 ± 3.36<br>*67.68 ± 1.22* | 222.62 ± 6.10<br>*73.09 ± 1.14* | 227.88 ± 1.48<br>*73.09 ± 1.14* |
| **$A_{iso}$ (nm)** | 114.71 ± 1.62<br>*68.70 ± 1.00* | 103.77 ± 4.40<br>*68.11 ± 1.18* | 98.27 ± 3.28<br>*69.34 ± 1.20* | 95.11 ± 0.69<br>*69.34 ± 1.20* |
| **$A_{roti}$ (nm)** | -11.89 ± 1.84<br>*-2.18 ± 0.85* | -24.47 ± 0.90<br>*-1.11 ± 1.20* | -20.43 ± 1.54<br>*-0.72 ± 1.27* | -45.57 ± 2.25<br>*-0.72 ± 1.27* |

**Table S2.** Elastic constants for the DX core.

| | n = 18 | n = 20 | n = 21A | n = 21B | n = 22 |
|---|---|---|---|---|---|
| **Y (pN)** | 3478.58 ± 23.56 | 2217.67 ± 15.33 | 2341.52 ± 17.48 | 2612.11 ± 11.72 | 2788.19 ± 21.61 |
| **C (nm)** | 298.68 ± 5.02 | 221.59 ± 2.48 | 225.50 ± 8.48 | 226.66 ± 0.84 | 226.90 ± 2.88 |
| **k (nm/turn)** | 0.31 ± 0.02 | 0.67 ± 0.02 | 0.77 ± 0.07 | 0.57 ± 0.01 | 0.79 ± 0.04 |
| **$A_{ro}$ (nm)** | 139.30 ± 3.05 | 143.88 ± 4.19 | 151.04 ± 3.82 | 130.00 ± 0.32 | 131.93 ± 2.80 |
| **$A_{ti}$ (nm)** | 442.59 ± 0.84 | 615.95 ± 1.22 | 683.99 ± 3.25 | 672.13 ± 19.15 | 683.10 ± 4.32 |
| **$A_{iso}$ (nm)** | 211.89 ± 3.63 | 233.26 ± 5.63 | 245.62 ± 4.29 | 217.01 ± 0.85 | 220.61 ± 4.11 |
| **$A_{roti}$ (nm)** | 2.18 ± 0.72 | 1.39 ± 5.18 | 27.58 ± 4.70 | -18.56 ± 3.78 | -14.85 ± 5.68 |

### References


1. Beer, F.P., Johnston, E.R., DeWolf, J.T. and Mazurek, D.F. (2014) *Mechanics of Materials*. 7th ed. McGraw Hill.
2. Zhou, L., Marras, A.E., Su, H.-J. and Castro, C.E. (2014) DNA origami compliant nanostructures with tunable mechanical properties. *ACS Nano*, **8**, 27-34.
3. Lu, X.-J. and Olson, W.K. (2003) 3DNA: a software package for the analysis, rebuilding and visualization of three-dimensional nucleic acid structures. *Nucleic Acids Res.*, **31**, 5108-5121.
